# The Cardioprotective Effect of Ischemic Postconditioning is Mediated by Inhibiting RAP2C-MAP4K4 Pathway

**DOI:** 10.1101/2024.12.04.626922

**Authors:** Peng Yan, Zaixin Yu, Zhiqiang Hu, Sheng Li, Muka Mengjiang Juaiti, Min Zhang

## Abstract

**Background:** Ischemic postconditioning (PostC) serves as a vital defense for cardiomyocytes against the deleterious effects of ischemia/reperfusion (I/R) injury, the beneficial effects could be further enhanced through pharmacological strategies. Our prior research demonstrated upregulated expression of the GTP-binding protein RAP2C in H9C2 cells post hypoxia-reoxygenation (H/R). The cardioprotective effects of RAP2C and underlying mechanisms are unclear. We therefore explored the role of RAP2C in PostC-induced cardioprotection against I/R injury.

**Methods:** Open-chest rat I/R and primary cultured cardiomyocytes H/R models were used. RAP2C and MAP4K4 expression was detected by immunohistochemistry and Western blotting. The BioGRID and STRING databases were tapped to predict the RAP2C-MAP4K4 binding, which was confirmed by co-immunoprecipitation and immunofluorescence.

**Results:** Results indicated that I/R and H/R upregulated the protein levels of RAP2C, MAP4K4, phospho-JNK, phospho-P38, and phospho-ERK, concomitant with increased apoptosis. PostC mitigated these effects. The pro-apoptotic impacts and the activation of the MAPK pathway induced by H/R were attenuated by RAP2C knockdown and intensified by RAP2C overexpression. H/R increased the interaction between RAP2C and MAP4K4, and PostC attenuated this effect. MAP4K4 knockdown reduced the pro-apoptotic and MAPK-activating effects induced by both RAP2C overexpression and hypoxia/reoxygenation (H/R).

**Conclusions:** These results demonstrate that PostC reduces cardiomyocyte apoptosis via modulating RAP2C/MAP4K4 pathways, suggesting their potential as therapeutic targets for the treatment of ischemic heart disease.

## Introduction

Acute myocardial ischemia can cause irreversible cardiac damage and infarction, and reperfusion therapy is commonly used to alleviate myocardial ischemic injury ^1, 2^. However, this therapy can cause ischemia-reperfusion (I/R) injury (IRI), potentially leading to cell death and lethal myocardial reperfusion injury ^3, 4^. The occurrence of apoptosis in the early stage of reperfusion predicts the severity of IRI ^5^. Targeting apoptosis can prevent and attenuate IRI ^6^.

Ischemic postconditioning (PostC) is a therapeutic strategy referring brief intermittent cycles of ischemia and reperfusion immediately after an ischemic event ^7^. PostC reduces cardiomyocyte loss by inhibiting inflammation, apoptosis, ROS generation, and intracellular calcium overload^8, 9^. These processes involve multiple mediators, end effectors, and pathways, as well as changes in the mitochondrial permeability transition pore (mPTP) through a complex signal transduction cascade ^8, 10, 11^. However, the molecular mechanisms and pharmacological targets of PostC are largely unknown.

Ras-Associated Protein 2C (RAP2C) is a member of the Ras superfamily of GTPases. RAP2C has a lower binding affinity to GTP and a slower rate of release of GDP than other Ras proteins ^12–14^. As a molecular switch for signal transduction, RAP2C regulates mitochondrial fusion and metabolism as well as cell apoptosis, proliferation, and migration in tumors ^15–17^. We previously showed that RAP2C was linked with enhanced apoptosis in H9C2 cells post hypoxia/reoxygenation (H/R)^18^. However, little is known about the roles of RAP2C in cardiovascular function. Moreover, the mechanisms by which RAP2C promotes cardiomyocyte apoptosis during IRI and regulates downstream signaling are unclear.

The analysis of the BioGRID and STRING databases suggests that RAP2C interacts with mitogen-activated protein kinase kinase kinase kinase-4 (MAP4K4). MAP4K4 is an upstream kinase of the Mitogen-Activated Protein Kinase (MAPK) cascade and plays a central role in cardiomyocyte death. In cultured rat cardiomyocytes, rodent models of myocardial IRI, and human heart tissue, IRI could activate MAP4K4, which upregulates apoptotic proteins, including caspase-3 and TAK1 ^19, 20^. Traf2 and Nck-interacting kinase (TNIK), which share 90% amino acid identity with MAP4K4, binds to RAP2 ^21^, triggering RAP2 autophosphorylation and JNK-MAPK pathway activation, suggesting that MAP4K4 interacts with RAP2 ^22^. MAP4K4 activates the MAPK pathway, which governs stress responses and cell fate, proliferation, and migration ^23^. Furthermore, the ERK1/2 pathway inhibitor PD98059 reversed the effect of PostC on infarct size and left ventricular cardiomyocyte loss ^24^. Therefore, it is reasonable to speculate that RAP2C might induce myocardial apoptosis by activating MAPK signaling via MAP4K4.

This study assessed the roles of RAP2C in PostC-mediated cardioprotection against IRI and underlying mechanisms using *in vitro* assays and a rat model of myocardial IRI. We hypothesized that RAP2C downregulation by PostC decreased H/R injury (HRI) in cardiomyocytes by suppressing the MAPK pathway through MAP4K4.

## Materials and methods

All experimental procedures were performed in accordance with the guidelines established by the European Ethical Committee (EEC) (2010/63/EU) and the National Institutes of Health Guide for the Care and Use of Laboratory Animals. The protocols were approved by the Animal Care and Use Committee of Central South University (CSU-2023-0169).

### Primary cardiomyocyte culture, transfection, and cellular model of HRI

Primary cardiomyocytes were isolated from the hearts of 1-2 day-old Sprague-Dawley rats, as described previously ^25^ (Online Supplementary Material, Video S1). Cells were transfected with siRNA (50 nM, RiboBio, China) using the riboFECT CP reagent (RiboBio, China) for 24 h, according to the manufacturer’s instructions. The siRNA sequences of RAP2C and MAP4K4 are shown in Table S1. Then, cells were transfected with a RAP2C-overexpressing adenovirus (Ad-mCherry-RAP2C, Hanbio, China) or a control adenovirus at a multiplicity of infection of 50. Cardiomyocytes were randomly allocated to three groups: normoxia (N), H/R, and H/R plus PostC. In the H/R group, cardiomyocytes were cultured in Hank’s balanced salt solution in an anaerobic chamber (N_2_/CO_2_/H_2_: 85%/5%/10%) for 4 h. Cells were reoxygenated in fresh medium in a humidified incubator with 5% CO_2_ for 6 h. In the H/R plus PostC group, cells were cultured under the following conditions: a hypoxic environment (anaerobic chamber) for 4 h, a normoxic environment (humidified incubator with 5% CO_2_) for 5 min, followed by a hypoxic environment for 5 min. This cycle was repeated three times. Then, cells were reoxygenated in fresh medium in a humidified incubator with 5% CO_2_ for 6 h. In the N group, cells were cultured in a humidified incubator with 5% CO_2_ for 9 h ^8^. All incubations were performed at 37°C.

### Establishment of a rat model of IRI and measurement of infarct size

Male Sprague-Dawley rats (200-230 g) were anesthetized by the intraperitoneal injection of 25% urethane (4 mL/kg). Echocardiographic data were obtained and monitored using a BL410 Biological Function Experimental System (Techman Instrument Co., Ltd., Chengdu, China). As previously described^26^, small incision was made in the trachea, and underlying muscles were retracted. The trachea was cannulated, and the animals were connected to a small animal ventilator (HX-101E, Techman Instrument Co. Ltd. Chengdu, China). The tidal volume and respiration rate were set to 7.0 and 5:4. The cartilages of the third to fifth ribs were cut, and underlying muscles were retracted laterally. Then, we passed a 3-0 suture between the upper and lower ribs on each side, cut the ribs in the middle, and pulled the suture to opposite sides to separate the ribs. Once tightened and secured, interrupt the ribs in the middle. The pericardium was incised to expose the heart. A 7-0 suture was passed under the left anterior descending (LAD) coronary artery, and blood flow was controlled with a polyethylene tube and vascular clamps. The murine model of myocardial IRI was established as follows: ligation of the LAD artery for 30 min, followed by three 30 s cycles of reperfusion and ligation, and one 120 min cycle of reperfusion. The artery was ligated with a 7-0 silk suture approximately 3 mm below the anterior-inferior edge of the left atrium. The ends of the suture were threaded through a 1 cm polyethylene tube, which was placed parallel to the LAD artery to form a snare for reversible occlusion. After reperfusion for 120 min, the artery was ligated, and the area at risk was evaluated by injecting 1% Evans blue dye solution into the jugular vein. Local ischemia was confirmed by observing changes in echocardiographic parameters (ST elevation) and in the color of the myocardium. After exposing the heart, the left ventricle was removed, frozen at −20 °C for 30 min, and cut into five to six 0.5-mm sections from top to bottom. The sections were incubated with 1% 2,3,5-triphenyltetrazolium chloride (TTC) at 37°C for 15 min protected from light and fixed in 4% paraformaldehyde in PBS for 30 min. Images were analyzed using Image-Pro Plus 6.0 (Media Cybernetics, Bethesda, MD, USA).

### Western blotting

Proteins were extracted using RIPA buffer containing protease and phosphatase inhibitors (EpiZyme). Protein concentration was determined using a BCA Protein Assay kit. Proteins (10-20 μg) were separated by SDS-PAGE and transferred to polyvinylidene difluoride membranes. Membranes were blocked with 5% bovine serum albumin (BSA) in PBS for 1 h. The samples were probed with primary antibodies overnight at 4°C, followed by incubation with horseradish peroxidase (HRP)-conjugated secondary antibodies at room temperature. Immunoreactive bands were visualized using chemiluminescent reagents (Beyotime, China) and an enhanced chemiluminescence detection system (Bio-Rad, China). Fluorescence intensity was quantified using ImageJ 6.0 software ^27^. The primary and secondary antibodies are listed in Table S1.

### Flow cytometry

Apoptosis was measured using the Annexin V-FITC Apoptosis Detection Kit (Vazyme). Cardiomyocytes were washed twice with PBS and resuspended in 100 µL of 1× Annexin V binding buffer containing 5 µL of Annexin V-FITC and 5 µL of propidium iodide or 7-amino-actinomycin D. After incubation for 10 min protected from light, cells were resuspended in PBS and analyzed using a flow cytometer (DxP Athena, Cytek Biosciences, Fremont, CA, USA).

### Terminal deoxynucleotidyl transferase-mediated dUTP nick end-labeling (TUNEL) assay

The detection of apoptotic cells was performed using the TUNEL BrightGreen and BrightRed Apoptosis Detection Kit (Vazyme A113-02) following the manufacturer’s instructions. Green or Red is for apoptotic cells, and blue is for nuclei marked with DAPI. Cardiac tissues and cardiomyocytes were observed under a fluorescence microscope (Leica DMi8, Germany).

### Immunohistochemistry (IHC)

Cardiac tissues were fixed in 4% paraformaldehyde, embedded in paraffin, and cut into 5 µm sections. Tissue sections were deparaffinized in xylene, rehydrated in graded ethanol (100% twice, 90% once, 80% once, and 70% once, each for 5 min), and washed twice in PBS (5 min each). Antigen retrieval was performed by microwave heating in 0.01 M citrate buffer (pH 6.0) for 20 min. Endogenous peroxidase activity was blocked by incubating the sections in 1% periodic acid for 10 min at room temperature. Samples were probed with primary antibodies overnight at 4°C, rinsed with PBS, and incubated with HRP-conjugated goat anti-rabbit secondary antibody for 30 min at room temperature. Sections were stained with 3,3′-diaminobenzidine (DAB) and hematoxylin and rinsed with distilled water and PBS. Slides were dehydrated in a graded ethanol series (60-100%, 5 min each) and xylene (10 min, twice), sealed with a neutral gum and observed under a fluorescence microscope (Leica DMi8, Germany).

### Immunofluorescence

Myocardial tissue was fixed in 4% paraformaldehyde for 10 min and permeabilized in Triton X-100 1% for 20 min. Endogenous peroxidase activity was blocked using 5% BSA for 1 h. Tissues were incubated with primary antibodies against RAP2C (orb474405, Biorbyt, Cambridge, UK) and MAP4K4 (sc-100445, Santa Cruz) overnight at 4℃, followed by incubation with CoraLite594-conjugated goat anti-mouse antibody (SA00013-3, Proteintech) and CoraLite488-conjugated goat anti-rabbit antibody (SA00013-2, Proteintech) at room temperature for 1 h. Nuclei were stained with DAPI. The samples were washed thrice in PBS between each step and imaged on a confocal microscope (Leica SP8, Germany).

### Co-immunoprecipitation assay

Cardiomyocytes were infected with Ad-mCherry-RAP2C for 48 h and lysed with NP-40 for 30 min. The lysate was centrifuged at 12000*g* for 15 min at 4°C. The supernatant was incubated with a monoclonal antibody against RAP2C or rabbit IgG (control) overnight at 4°C, followed by incubation with protein A/G agarose beads for 1 h at 4°C. The beads were collected by centrifugation at 3000 rpm for 3 min and washed thrice with NP-40 lysis buffer. The immunocomplexes were resolved by SDS-PAGE, transferred to polyvinylidene difluoride membranes, and probed with a primary antibody against MAP4K4.

### Statistical analysis

Data were analyzed using GraphPad Prism version 8 and were expressed as mean ± standard deviation of at least three biological replicates. The three groups were compared using one-way analysis of variance followed by the least significant difference test. A *p*-value of less than 0.05 was considered statistically significant.

## Results

### *RAP2C* downregulation is associated with PostC-mediated cardioprotection

To study the regulation of RAP2C in PostC, we used rat model of ischemia/reperfusion injury (IRI) and cellular model of primary rat cardiomyocytes. The rat model of myocardial IRI exhibited characteristic ST-segment elevation after LAD coronary artery ligation in vivo (Fig. S1A), increased number of TUNEL positive cells, whereas PostC could reduce IRI (Fig.1A, B). Immunohistochemistry and Western blotting showed that I/R significantly increased myocardial RAP2C expression, while PostC attenuated these effects (Fig. 1C, D). In our cellular model of H/R (4 h hypoxia and 6 h reoxygenation), the apoptosis rate in cardiomyocytes increased significantly, without cell shedding or loss (Fig. S1B), which could be reduced by PostC (Fig. 1E). Western blots showed that H/R increased the Bax/Bcl-2 ratio and the expression of cleaved caspase 3, cleaved caspase 9, and RAP2C in cardiomyocytes, whereas PostC reversed these effects (Fig. 1F, G). These findings suggest that PostC reduces H/R-induced cardiomyocyte apoptosis and possibly by inhibiting RAP2C.

**Figure 1.**
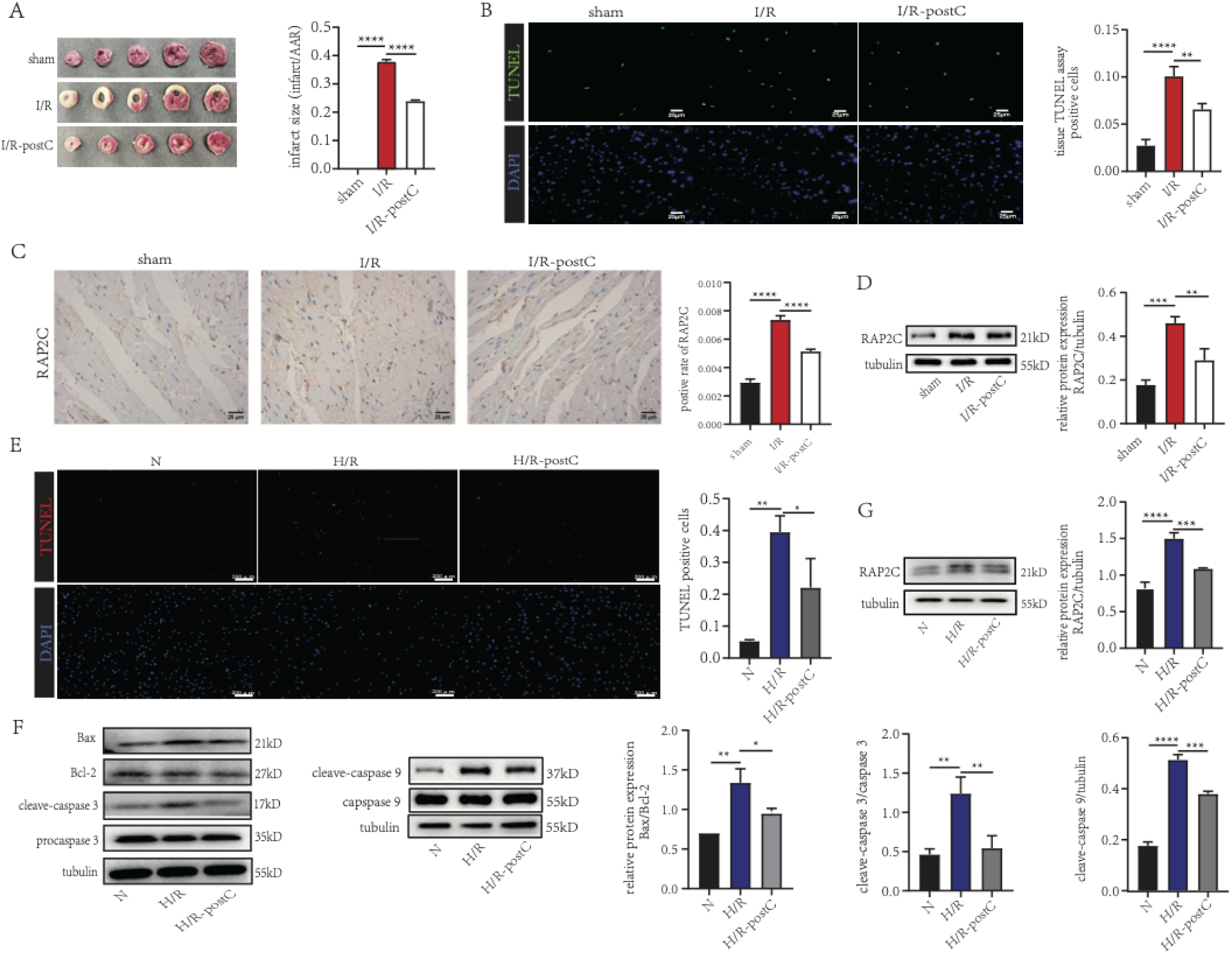
The cardioprotective effect of ischemic postconditioning (PostC) is mediated by RAP2C expression. **a.** Infarct size in rats subjected to sham treatment, ischemia/reperfusion (I/R), or I/R + PostC based on staining with 2,3,5-triphenyltetrazolium chloride. **b.** Number of TUNEL-positive cardiomyocytes in the study groups (×10) (scale bar = 25 µm). **c.** Immunohistochemical analysis of RAP2C expression in rat myocardial tissue based on the intensity of positive staining (integrated optical density per stained area) (scale bar = 25 µm). **d.** Western blot analysis of RAP2C expression in rat myocardial tissue. **e.** Number of TUNEL-positive cardiomyocytes in animals exposed to normoxia (N), hypoxia/reperfusion (H/R), or H/R + PostC (×10) (scale bar = 200 µm). **f, g.** Western blot analysis of the Bax/Bcl-2 ratio and the expression of cleaved caspase-3, cleaved caspase-9, and RAP2C in cultured cardiomyocytes. Data are means and standard deviations of three biological replicates. *p* values were calculated by two-tailed Student’s *t-*test. one-way ANOVA with Tukey’s post hoc test was used for multiple group comparisons. \**P* < 0.05, \*\**P* < 0.01, \*\*\**P* < 0.001, \*\*\*\**P* < 0.0001.

### PostC reduces H/R-induced cardiomyocyte apoptosis in a RAP2C-dependent manner

The role of RAP2C in rat cardiomyocytes exposed to H/R and H/R plus PostC was then evaluated by loss-of-function assays using small interfering RNA (siRNA) targeting *RAP2C*. The efficiency of the siRNA-mediated knockdown of *RAP2C* was assessed by Western blotting (Fig. 2A). Cardiomyocytes were transfected with siRAP2C or siNC and exposed to H/R or PostC after H/R. H/R increased the relative protein expression of RAP2C; PostC suppressed this effect and acted synergistically with siRAP2C (Fig. 2B). Similar results were obtained for the rate of apoptosis (Fig. 2C) and for Bax/Bcl-2 and cleaved caspase 9 expression in cardiomyocytes (Fig. 2D). Flow cytometry results showed that H/R significantly increased late apoptosis rate, whereas PostC attenuated this effect, which was further enhanced by *RAP2C* silencing (Fig. 2E). These results indicate that RAP2C downregulation contributes to the cardioprotective effect of PostC. To further validate the causal role of RAP2C on PostC mediated protective effects, *RAP2C* was overexpressed using an adenovirus system. Western blot analysis showed that transfection efficiency was satisfactory at a multiplicity of infection of 30 (Figs. S1C and 3A). Ad-RAP2C markedly enhanced the stimulatory effect of H/R on RAP2C expression, and blocked the downregulating effects of PostC postH/R (Fig. 3B). TUNEL staining showed similar results (Fig. 3C). Furthermore, *RAP2C* overexpression enhanced the stimulatory effect of H/R on Bax/Bcl-2, cleaved caspase-3, and cleaved caspase-9 expression, while the protective effects of PostC were blocked by Ad-RAP2C (Fig. 3D). These findings suggest that RAP2C enhances H/R induced cardiomyocyte apoptosis and PostC reduces this effect in a RAP2C-dependent manner.

**Figure 2.**
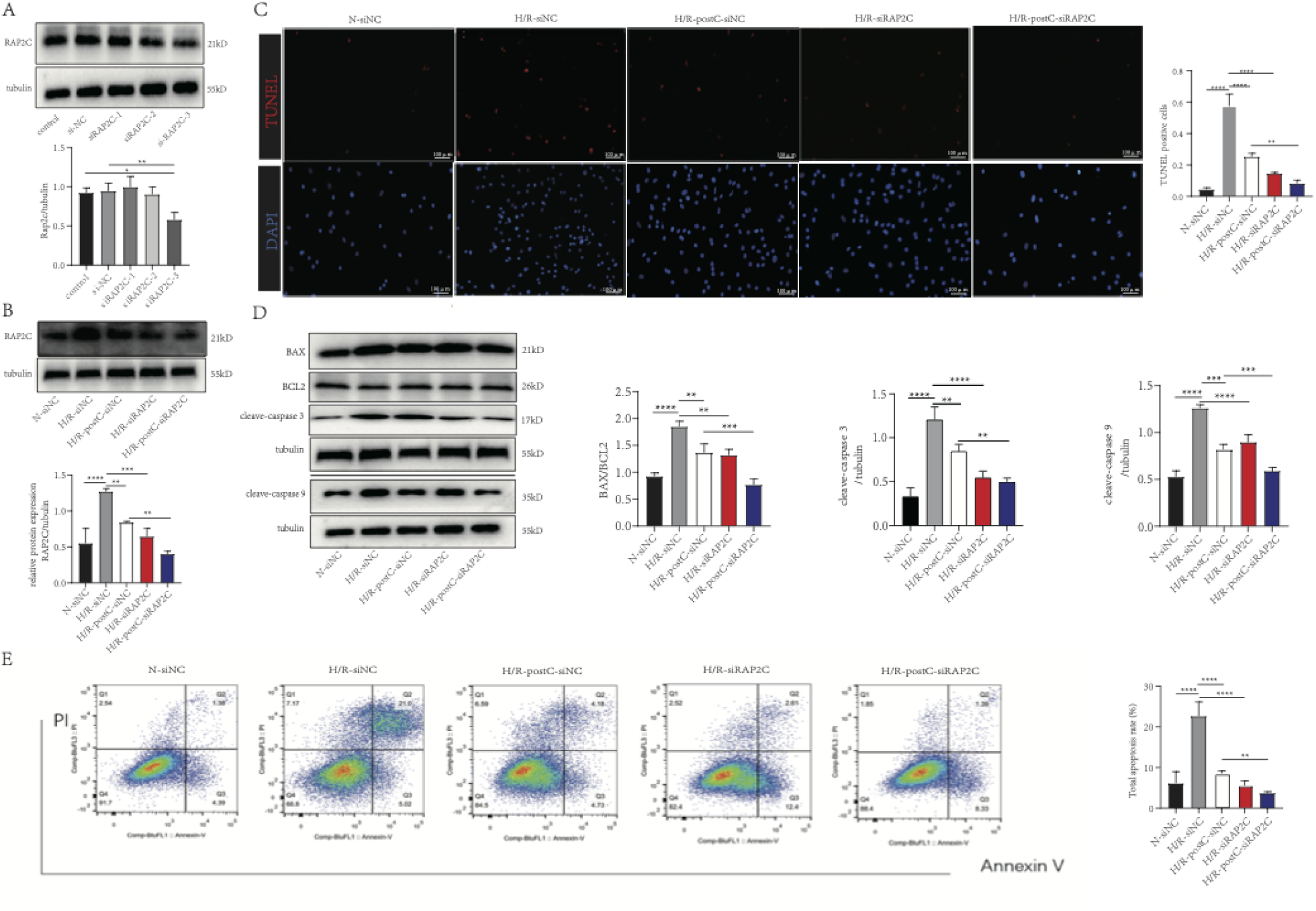
*RAP2C* silencing enhances the anti-apoptotic effect of ischemic postconditioning (PostC) in cardiomyocytes under hypoxia/reoxygenation (H/R). **a.** Western blot analysis of the efficiency of siRNA-mediated silencing of *RAP2C*. **b, d.** Western blot analysis of RAP2C expression in cardiomyocytes transfected with si-RAP2C or si-NC and exposed to normoxia (N), H/R, or H/R + PostC. **c.** Number of TUNEL-positive cells in different groups (×20) (scale bar = 100 µm). **d.** Western blot analysis of the Bax/Bcl-2 ratio and the expression of cleaved caspase-3 and −9. **e**. Flow cytometry analysis of total and late apoptosis rates. Data are means and standard deviations of three biological replicates. *p* values were calculated by two-tailed Student’s *t-*test. one-way ANOVA with Tukey’s post hoc test was used for multiple group comparisons. \**P* < 0.05, \*\**P* < 0.01, \*\*\**P* < 0.001, \*\*\*\**P* < 0.0001.

**Figure 3.**
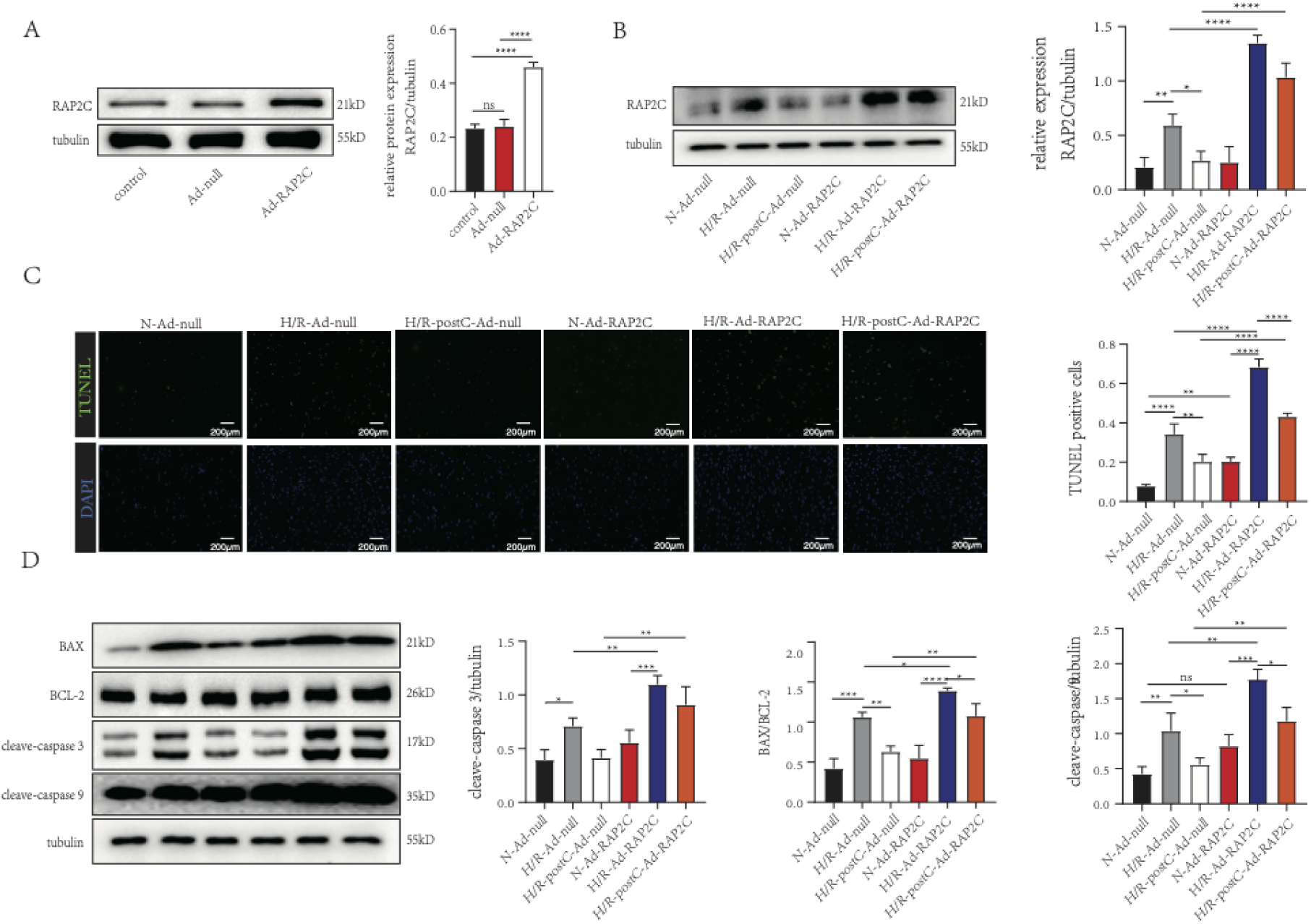
RAP2C enhances hypoxia/reoxygenation (H/R)-induced cardiomyocyte apoptosis. **a.** Western blot analysis of the efficiency of *RAP2C* overexpression using an adenovirus system. **b.** Western blot analysis of RAP2C expression in cardiomyocytes infected with Ad-null or Ad-RAP2C and exposed to normoxia, H/R, or H/R + ischemic postconditioning (PostC). **c.** Number of TUNEL-positive cells in different groups (scale bar = 200 µm). **d.** Western blots of the Bax/Bcl-2 ratio and the expression of cleaved caspase-3 and −9. Data are means and standard deviations of three biological replicates. *p* values were calculated by two-tailed Student’s *t-*test. one-way ANOVA with Tukey’s post hoc test was used for multiple group comparisons. \**P* < 0.05, \*\**P* < 0.01, \*\*\**P* < 0.001, \*\*\*\**P* < 0.0001.

### PostC mitigates cardiomyocyte apoptosis by inhibiting the RAP2C-MAPK pathway

The MAPK signaling pathway is involved in cardiomyocyte apoptosis, oxidative stress, and inflammation under H/R conditions ^28, 29^. Thus, we hypothesized that RAP2C increased cardiomyocyte apoptosis by regulating this pathway. Western blot analyses of myocardial tissue and cardiomyocytes showed that I/R and H/R increased MAPK phosphorylation compared with normoxia controls. The MAPK family, known for its key regulatory roles, comprises members such as Extracellular signal-regulated kinase (ERK), Jun N-terminal kinase (JNK), and p38. Notably, PostC decreased this effect (Fig. 4A, B). H/R significantly increased the expression of MAPK compared to normoxia, whereas *RAP2C* silencing and PostC reduced these effects (Fig. 4C). H/R also upregulated these MAPK components in Ad-null infected mice, while PostC abrogated this effect. *RAP2C* overexpression blocked the beneficial effects of PostC (Fig. 4D this important to show the causal relationship, pls modify 4D as suggested). These findings demonstrate that the MAPK pathway is involved in RAP2C-induced cardiomyocyte apoptosis under H/R conditions.

**Figure 4.**
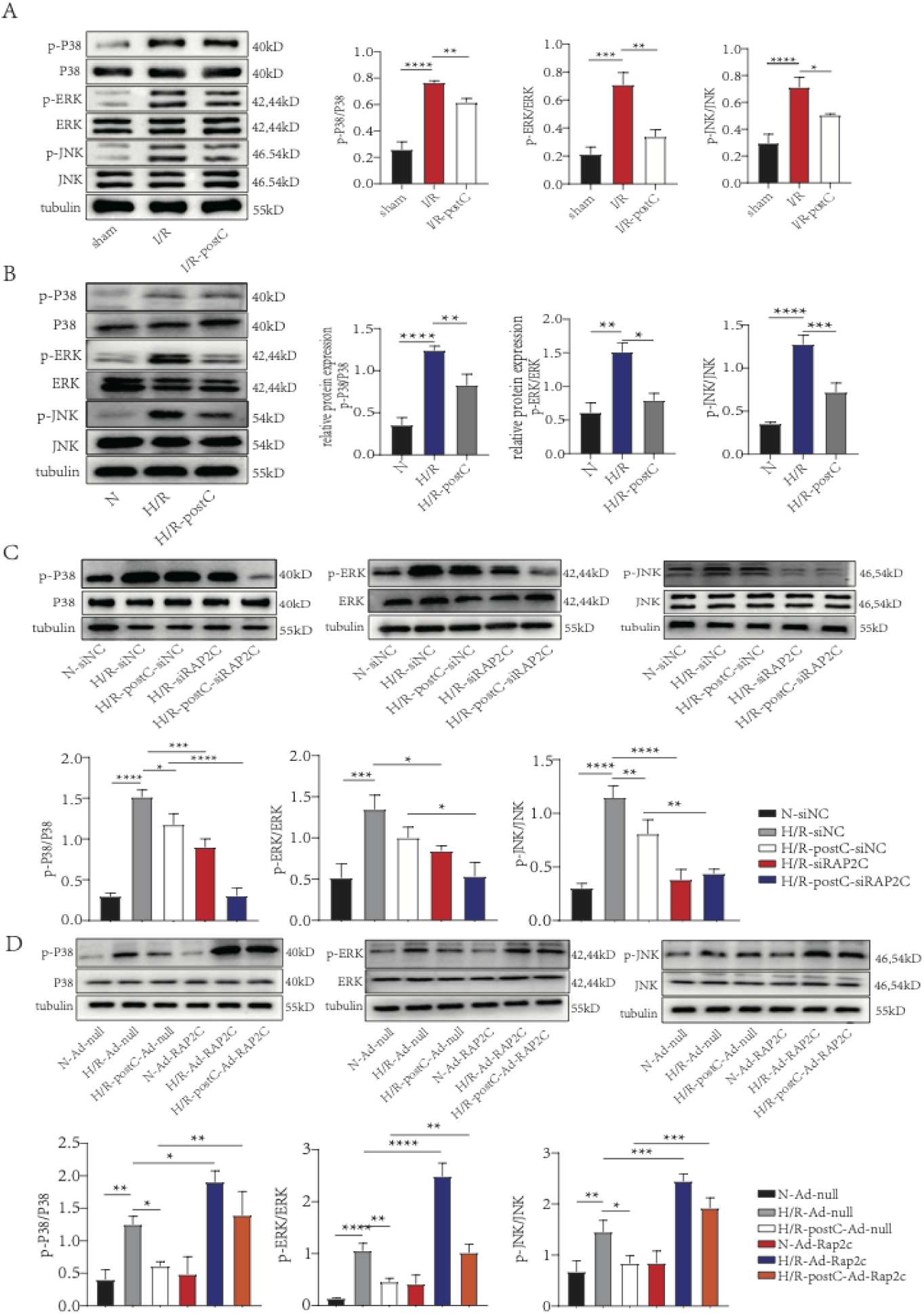
Ischemic postconditioning (PostC) mitigates cardiomyocyte apoptosis by inhibiting the RAP2C-MAPK pathway. **a.** Western blot analysis of the relative expression of MAPK components in rat myocardial tissue exposed to sham conditions, ischemia/reperfusion (I/R), or I/R + PostC. **b.** Western blots of the relative protein expression of MAPK components in cardiomyocytes exposed to normoxia, hypoxia/reperfusion (H/R), or H/R + PostC. **c.** MAPK expression in cardiomyocytes transfected with si-RAP2C or si-NC and exposed to normoxia (N), H/R, or H/R + PostC. **d.** MAPK expression in cardiomyocytes transduced with Ad-RAP2C or Ad-null and exposed to normoxia (N), H/R, or H/R + PostC. Data are means and standard deviations of three biological replicates. *p* values were calculated by two-tailed Student’s *t-*test. one-way ANOVA with Tukey’s post hoc test was used for multiple group comparisons. \**P* < 0.05, \*\**P* < 0.01, \*\*\**P* < 0.001, \*\*\*\**P* < 0.0001.

### MAP4K4 participates in PostC-mediated cardioprotection against IRI and H/R

The analysis of the BioGRID and STRING databases suggests that MAP4K4 binds to RAP2C (Fig. 5A). IHC and western blots showed that I/R increased MAP4K4 expression compared with sham conditions, while PostC suppressed this effect (Fig. 5B, C). Similar results were observed in cardiomyocytes under H/R conditions (Fig. 5D), and *RAP2C* knockdown and PostC reduced this increase (Fig. 5E). Consistent with this finding, *RAP2C* overexpression enhanced the upregulation of MAP4K4 by H/R, and reversed the beneficial effects of PostC (Fig. 5F.pls modify this figure as suggested to show the causal relationship), indicating that RAP2C acted as a positive regulator of MAP4K4. Co-immunoprecipitation assays suggest that MAP4K4 binds to RAP2C in cardiomyocytes under normoxia. H/R enhanced this interaction, while PostC attenuated this effect (Fig. 5G). Immunofluorescence showed weak colocalization of RAP2C and MAP4K4 around the nucleus in normoxic cardiomyocytes. This result is consistent with the quantification of colocalization using Pearson’s correlation coefficient (Fig. 5H). These data indicate that PostC reduces myocardial apoptosis by weakening the interaction between RAP2C and MAP4K4.

**Figure 5.**
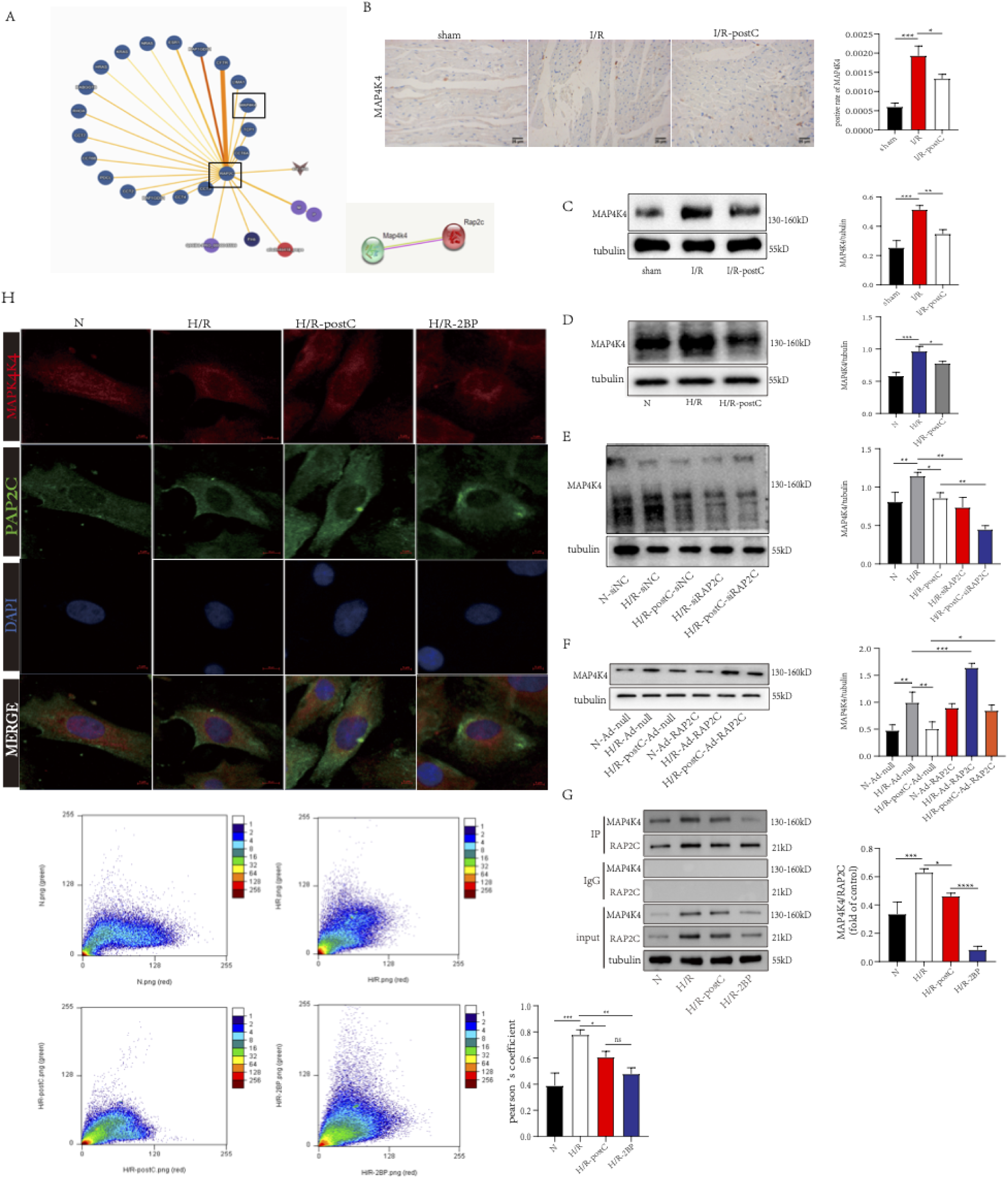
Ischemic postconditioning (PostC) alleviates cardiomyocyte apoptosis by inhibiting the RAP2C-MAP4K4 pathway. **a.** Protein interactions based on the analysis of the BioGRID and STRING databases. **b.** Immunohistochemical analysis of MAP4K4 expression in rat myocardial tissue exposed to sham, I/R and I/R-postC in rat myocardial tissue (scale bar = 25 µm). **c.** MAP4K4 expression in rat cardiomyocytes exposed to sham conditions, ischemia/reperfusion (I/R), or I/R + PostC. **d.** MAP4K4 expression in rat cardiomyocytes exposed to normoxia (N), hypoxia/reoxygenation (H/R), or H/R + PostC. **e.** MAP4K4 expression in cardiomyocytes transfected with si-RAP2C or si-NC and exposed to H/R or H/R + PostC. **f.** MAP4K4 expression in cardiomyocytes transduced with Ad-RAP2C or Ad-null and exposed to H/R or H/R + PostC. **g.** Co-immunoprecipitation assays using cardiomyocyte lysates and RAP2C antibody under normoxia, H/R, and H/R + PostC. MAP4K4 and RAP2C expression in input and immunoprecipitated samples was evaluated by western blotting. **h.** Colocalization of MAP4K4 and RAP2C by immunofluorescence double staining (scale bar = 5/10 μm). The degree of colocalization is shown in scatter plots and was calculated using Pearson’s correlation coefficient. Data are means and standard deviations of three biological replicates. *p* values were calculated by two-tailed Student’s *t-*test. one-way ANOVA with Tukey’s post hoc test was used for multiple group comparisons. \**P* < 0.05, \*\**P* < 0.01, \*\*\**P* < 0.001, \*\*\*\**P* < 0.0001.

### MAP4K4 mediates the anti-apoptotic effect of PostC

We then determined whether the MAP4K4-MAPK pathway was involved in the anti-apoptotic effect of PostC. MAP4K4 expression was significantly lower in cardiomyocytes transfected with siMAP4K4 than in mock-transfected cells (Fig. 6A). H/R increased the expression of p-ERK, p-JNK, and p-P38 in cardiomyocytes, and PostC and siMAP4K4 abrogated this effect (Fig. 6B). H/R increased the number of TUNEL positive cardiomyocytes and total apoptosis rate, while PostC and siMAP4K4 diminished this effect (Fig. 6C, D). Further, H/R significantly increased the expression of cleaved caspase 3 and caspase 9 and the Bax/Bcl-2 ratio, whereas PostC and siMAP4K4 reduced this effect (Fig. 6E). These results indicate that the anti-apoptotic effect of PostC is possibly mediated by the inhibition of the MAP4K4-MAPK pathway. Here we need to show the beneficial effects of PostC could be blocked by OE(Ad) MAP4K4, without this, we can only say “possible” not causal.

**Figure 6.**
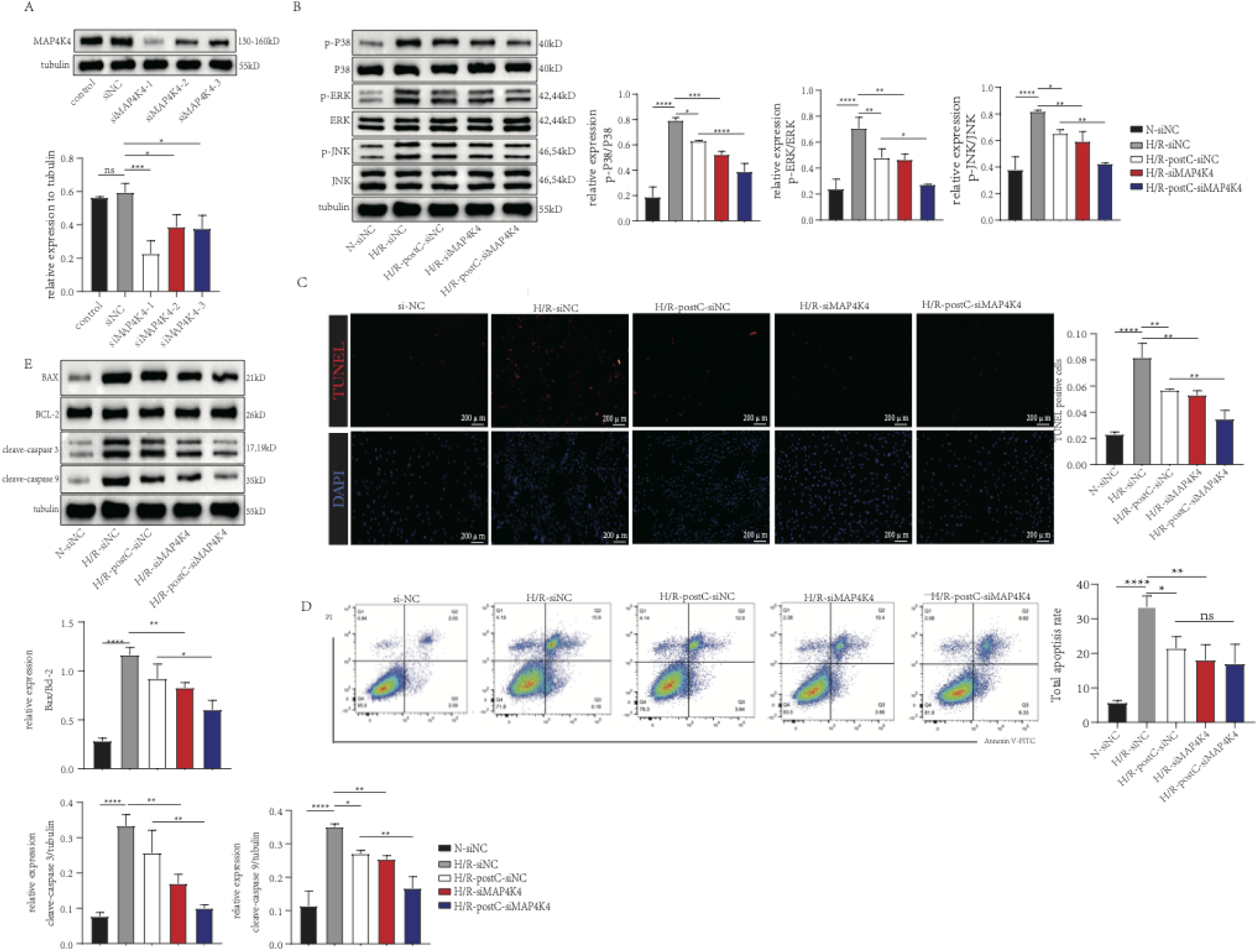
si-MAP4K4 enhances the anti-apoptotic effect of ischemic postconditioning (PostC) in rat cardiomyocytes exposed to hypoxia/reoxygenation (H/R). **a.** Western blot analysis of the efficiency of siRNA-mediated silencing of MAP4K4 expression. **b.** p-ERK, p-JNK, and p-P38 expression in cardiomyocytes transfected with si-MAP4K4 or si-NC and exposed to normoxia (N), H/R, or H/R + PostC. **c.** Number of TUNEL-positive cells in different groups (×10) (scale bar = 200 µm). **d.** Flow cytometry analysis of total apoptosis rates in different groups. **e.** Western blot analysis of the expression of cleaved caspase-3 and −9 and the Bax/Bcl-2 ratio. Data are means and standard deviations of three biological replicates. \**P* < 0.05, \*\**P* < 0.01, \*\*\**P* < 0.001, \*\*\*\**P* < 0.0001.

### SiMAP4K4 reduces the pro-apoptotic effect of Ad-RAP2C in cardiomyocytes

To confirm that RAP2C aggravates apoptosis via MAP4K4, we measured the expression of MAP4K4 and RAP2C in rat cardiomyocytes transfected with Ad-RAP2C or Ad-RAP2C combined with siMAP4K4 and exposed to H/R and PostC. As expected, Ad-RAP2C increased RAP2C and MAP4K4 expression; siMAP4K4 reduced the effect of Ad-RAP2C on MAP4K4 expression but not on RAP2C expression (Fig. 7A), indicating that MAP4K4 is the downstream of RAP2C. Furthermore, siMAP4K4 reduced the Ad-RAP2C-induced upregulation of p-ERK, p-JNK, and p-P38 (Fig. 7B). TUNEL staining, flow cytometry, and Western blots revealed that *MAP4K4* knockdown eliminated the pro-apoptotic effect of RAP2C in H/R cardiomyocytes (Fig. 7C-E). These data suggested that MAP4K4 is actively participated on RAP2C-induced myocardial apoptosis. Another Good-way is to block the PostC effects by ad-MAP4K4.

**Figure 7.**
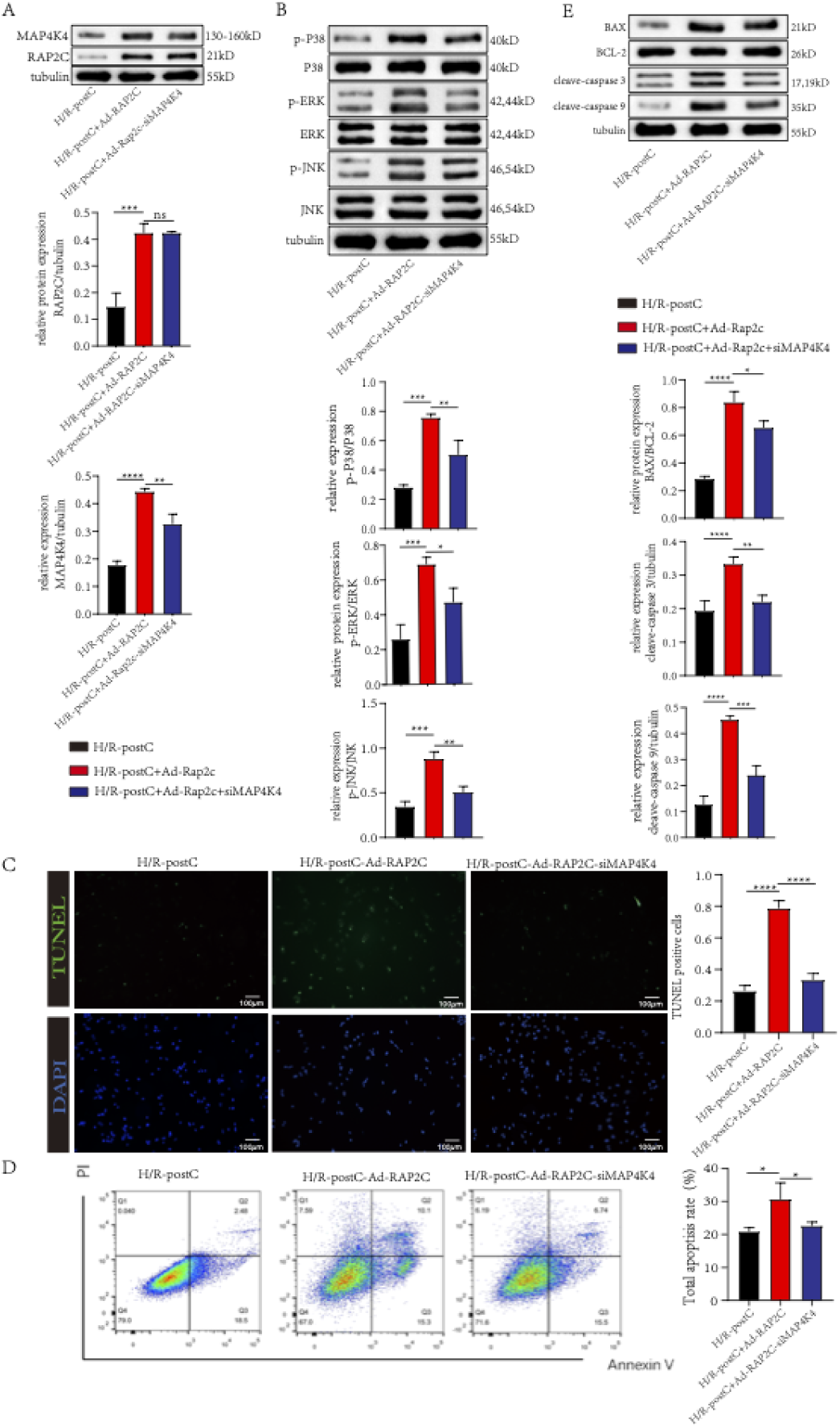
si-MAP4K4 reduces the pro-apoptotic effect of Ad-RAP2C in rat cardiomyocytes. **a.** Western blot analysis of RAP2C and MAP4K4 expression in cells transfected with Ad-RAP2C or Ad-RAP2C + si-MAP4K4 and exposed to hypoxia/reoxygenation (H/R) and ischemic postconditioning (PostC). **b.** Protein expression of p-ERK, p-JNK, and p-P38 in different groups. **c.** Number of TUNEL-positive cells in different groups (×10) (scale bar = 100 µm). **d.** Flow cytometry analysis of the rate of apoptosis. **e.** Western blot analysis of the Bax/Bcl-2 ratio and the expression of cleaved caspase-3 and −9 in cardiomyocytes. Data are means and standard deviations of three biological replicates. *p* values were calculated by two-tailed Student’s *t-*test. one-way ANOVA with Tukey’s post hoc test was used for multiple group comparisons. \**P* < 0.05, \*\**P* < 0.01, \*\*\**P* < 0.001, \*\*\*\**P* < 0.0001.

## Discussion

Present study showed that the anti-apoptotic effect of PostC was mediated by RAP2C and MAP4K4 downregulation. To our best knowledge, this is the first report describing the RAP2C and MAP4K4 related protective mechanism of PostC in IRI and H/R models.

RAP2C is a RAS GTPase with 91.3% homology with human RAP2A. RAP2C has three functional domains: an N-terminal domain that interacts with downstream effector proteins, a GTPase domain, and a domain containing a CAAX motif that targets Ras proteins to the plasma membrane. RAP2C participates in multiple processes, including cellular metabolism, tumor progression, and nervous system structure and function ^15, 16, 30–33^. However, the roles of RAP2C in cardiovascular function are poorly understood. The results of TUNEL staining and flow cytometry showed that *RAP2C* silencing enhanced the anti-apoptotic effect of PostC, whereas *RAP2C* overexpression attenuated the protective effect and increased IRI, demonstrating that RAP2C worsened IRI and that PostC reduced cardiomyocyte apoptosis by downregulating RAP2C.

Apoptosis is a crucial aspect of the pathogenesis of myocardial IRI ^34, 35^. *RAP2C* is upregulated in tumors and may function as an oncogene and anti-apoptotic factor in lung squamous cell carcinoma, small-cell lung cancer, hepatocellular carcinoma, and breast cancer ^16, 32^. We previously showed that RAP2C increased apoptosis and decreased the viability of H9C2 cells under H/R conditions ^18^, implying that RAP2C contributes to apoptosis. Hypoxia activates the intrinsic pathway, opening mPTPs and releasing Smac/DIABLO and Omi/HtrA2 into the cytoplasm. Apaf-1 is oligomerized in the presence of procaspase-9, dATP, and cytochrome c, forming an apoptosome. Subsequently, caspase-9 activates caspase-3/7, initiating a positive feedback loop ^36, 37^. Our findings showed that RAP2C induced cardiomyocyte apoptosis by upregulating cleaved caspase-3, cleaved caspase-9, and Bax, suggesting that PostC reduced apoptosis via the intrinsic pathway, not extrinsic or perforin pathways. However, the anti-apoptotic effects of RAP2C via the intrinsic pathway warrant further investigation. Moreover, we found that RAP2C induced apoptosis via MAPK signaling. The Ras-Raf-MEK-MAPK pathway regulates cellular differentiation, apoptosis, and proliferation ^38–41^. RAP2C may control the MAPK pathway, which regulates cardiomyocyte apoptosis ^42^. The microRNA 188-5p targets *RAP2C* and promotes apoptosis in breast cancer cells by activating the MAPK pathway ^16^. This pathway also exacerbates myocardial apoptosis during I/R ^43^. For instance, *GADD45A* silencing and Prdx1 reduced H/R-induced apoptosis by inhibiting the ROS-activated P38 and JNK pathways ^44^. NEAT1 exacerbated myocardial IRI by stimulating MAPK signaling ^45^. We found that *RAP2C* overexpression increased P38/ERK/JNK phosphorylation induced by H/R and H/R plus PostC. In contrast, *RAP2C* silencing downregulated p-P38, p-ERK, and p-JNK, suggesting that RAP2C increased cardiomyocyte apoptosis by activating MAPK signaling.

MAP4K4, a member of the Ste20 kinase superfamily, regulates the pathogenesis of cardiovascular diseases, including obesity, atherosclerosis, and IRI ^46, 47^. MAP4K4 activation is associated with apoptosis and chronic heart failure in animals and humans ^19^. However, the role of MAP4K4 in IRI is unclear. Additionally, MAP4K4 substrates and inhibitors are currently unavailable. MAP4K4 controls endothelial barrier function and permeability through Rap GTPases ^48^. MAP4K4 binds to RAP2 to activate the JNK pathway in NIH3T3 cells ^22^. RAP2 binds to the citron homology domain (CNH) of the GCK-IV kinases MAP4K4, MINK, and TNIK and regulates the actin cytoskeleton by activating MAPK signaling ^49^. RAP2A binds to the CNH region of MINK to stimulate its kinase activity and induce TANC1 phosphorylation in synapses ^50^. RAP2A and its interaction with TNIK in the Wnt/β-catenin pathway regulate the stability of the LRP6 receptor ^51^. GTP-bound RAP2A promotes the MAP4K4-induced activation of JNK ^22^. These GCK-IV kinases share 90% amino acid identity in the CNH region ^52^. Additionally, RAP2C has more than 90% sequence homology with RAP2A, indicating that they may interact with the same effector proteins ^14^. These results suggest that RAP2C binds to MAP4K4 and phosphorylates MAPK components in cardiomyocytes.

This hypothesis was confirmed by co-immunoprecipitation and immunofluorescence assays showing that RAP2C co-eluted and co-localized with MAP4K4. H/R increased the strength of binding, whereas PostC abrogated this effect. *RAP2C* knockdown and overexpression in cardiomyocytes confirmed that MAP4K4 was a downstream effector of RAP2C. These results indicate that H/R modulates the expression and activity of RAP2C and MAP4K4. The interaction of these proteins may impair energy metabolism and increase free radical production and myocardial apoptosis^53, 54^. MAP4K4 is highly expressed in human spermatogonial stem cells and regulates cell proliferation and apoptosis through autophosphorylation and JNK activation ^55^. Under low extracellular matrix stiffness, RAP2 activates MAP4K4, leading to the activation of LATS1 and YAP ^56^. Since many Ste20 kinases are phosphorylated at multiple sites by other kinases or by autophosphorylation ^57, 58^ and we found that H/R strengthened the RAP2C-MAP4K4 interaction, we hypothesize that RAP2C recruits MAP4K4 to enhance autophosphorylation. Nonetheless, further research is needed to identify the binding sites of RAP2C and MAP4K4 and to assess whether RAP2C affects the phosphorylation activity of MAP4K4. We found that MAP4K4 regulated the MAPK pathway and cardiomyocyte apoptosis since si-MAP4K4 decreased MAPK activity and reduced apoptosis. PostC combined with *RAP2C* or *MAP4K4* knockdown, but not PostC alone, increased myocardial salvage and decreased downstream signaling. Apoptosis and the expression of three main MAPK components were reduced by PostC after *MAP4K4* knockdown and *RAP2C* overexpression. These results indicate that *MAP4K4* knockdown reverses *RAP2C* overexpression-induced apoptosis and MAPK pathway activation in cardiomyocytes. We assessed the effect of RAP2C on MAP4K4 expression and showed that PostC downregulated RAP2C *in vivo* and *in vitro*. These results suggest that MAP4K4 is a downstream effector of RAP2C and phosphorylates MAPK proteins, ultimately reducing cardiomyocyte apoptosis. Nonetheless, the effects of I/R + PostC on non-apoptotic cell death pathways warrant further investigation ^59^.

In conclusion, we showed that RAP2C upregulation exacerbated cardiac IRI. PostC reduced cardiomyocyte apoptosis by inhibiting the RAP2C-MAP4K4 pathways. Thus, RAP2C and MAP4K4 might serve promising therapeutic targets for IRI.

## Conclusion

PostC reduces cardiomyocyte apoptosis via modulating RAP2C/MAP4K4 pathways, suggesting their potential as therapeutic targets for the treatment of ischemic heart disease.

## Declarations section

### Availability of data and Materials

The datasets used and/or analysed during the current study are available from the corresponding author upon reasonable request.

### Ethics Approval and Consent to participate

The study was reviewed and approved by the Animal Care and Use Committee of Central South University (CSU-2023-0169).

### Author’s Contributions

All authors contributed to the study conception and design. The manuscript was drafted by M.Z and P.Y. The data were collected and analyzed by S.L and M.M.J. The manuscript was corrected and supervised by Z.X.Y and Z.Q.H. All authors consent to being responsible for all facets of the research, ensuring that the accuracy and integrity of every aspect of the work are thoroughly examined and addressed. All authors have endorsed the final manuscript and agree to publish.

## Consent for publication

Not applicable.

## Abbreviations Definitions

PostC: postconditioning

I/R: ischemia/reperfusion

H/R: hypoxia-reoxygenation

HRI: H/R injury

IRI: ischemia-reperfusion injury

mPTP: mitochondrial permeability transition pore

RAP2C: Ras-Associated Protein 2C

MAP4K4: mitogen-activated protein kinase kinase kinase kinase-4

MAPK: Mitogen-Activated Protein Kinase

TNIK: Traf2 and Nck-interacting kinase

EEC: European Ethical Committee

LAD: left anterior descending

TTC: triphenyltetrazolium chloride

BSA: bovine serum albumin

HRP: horseradish peroxidase

TUNEL: transferase-mediated

dUTP: nick end-labeling

DAB: diaminobenzidine

siRNA: small interfering RNA

ERK: Extracellular signal-regulated kinase

JNK: Jun N-terminal kinase

CNH: citron homology domain

